# Graph Transformer for drug response prediction

**DOI:** 10.1101/2021.11.29.470386

**Authors:** Thang Chu, Tuan Nguyen

**Affiliations:** Faculty of Mathematical Economics, National Economics University, Hanoi, Vietnam; Faculty of Science, University of Alberta, Edmonton, Canada

**Author notes:** Corresponding author: Tuan Nguyen.

**Keywords:** Drug response prediction, Graph Transformer, Kernel PCA, Deep learning, Graph convolutional network, Saliency map

## Abstract

**Background:** Previous models have shown that learning drug features from their graph representation is more efficient than learning from their strings or numeric representations. Furthermore, integrating multi-omics data of cell lines increases the performance of drug response prediction. However, these models showed drawbacks in extracting drug features from graph representation and incorporating redundancy information from multi-omics data. This paper proposes a deep learning model, GraTransDRP, to better drug representation and reduce information redundancy. First, the Graph transformer was utilized to extract the drug representation more efficiently. Next, Convolutional neural networks were used to learn the mutation, meth, and transcriptomics features. However, the dimension of transcriptomics features is up to 17737. Therefore, KernelPCA was applied to transcriptomics features to reduce the dimension and transform them into a dense presentation before putting them through the CNN model. Finally, drug and omics features were combined to predict a response value by a fully connected network. Experimental results show that our model outperforms some state-of-the-art methods, including GraphDRP, GraOmicDRP.

**Availability of data and materials:** https://github.com/chuducthang77/GraTransDRP.

## 1 Introduction

PERSONALIZED medicine is a rapidly advancing field in finding the specific treatment best suited for an individual based on their biological characteristic. Its approach relies on the understanding of an individual’s molecular and genomics profile [1]. However, conducting drug trials on each individual to observe corresponding responses is expensive and not even ethical. These challenges continue to be problematic for large-scale researches on this topic [2]. Therefore, cell lines of tumor samples have been obtained and developed as “artificial patients” to study the drug response.

Recently, projects such as GDSC [3], CCLE [4] and NCI60 [5] have published a database containing different types of omics data, such as copy number aberration, gene expression, methylation, etc. These data help to facilitate the development of computational methods for drug response prediction [6], [7], [8], [9]. As a result, competitions (Dream challenge) have been opened to direct the attention of researchers, scientists, and scholars toward solving this problem [10]. From simple to complex models, various machine learning algorithms have been applied to solve the drug sensitivity problem, namely support vector machines (SVMs), linear regression, and neural networks models [11] [12] [13]. More advanced methods such as multiple-kernel, multiple-task learning, and collaborative filtering techniques were proposed to integrate various types of - omics data or integrate different individual models to boost the performance [14] [15], [16], [17]. Meanwhile, graphbased methods focused on biological perspective have been introduced, including structural similarity between drugs, the biological similarity between cell lines, the interaction between proteins and gene regulatory [18] [19].

The machine learning-based methods above have proven their ability through the accuracy of drug response prediction. However, limitations exist, like drugs and cell lines’ representation. High dimensional data often represent them since each -omics profile can contain thousands of genes for each cell line. Similarly, there are a large number of chemical and structural features for each drug. Due to the limited number of cell lines, current machine learning methods have to face with “small *n*, large *p*” or “The curse of dimensionality” problem. As the number of dimensions increases, the volume of our domain increases exponentially, and it requires more samples so that the model can learn efficiently. As a result, traditional machine learning-based methods will likely become underfitting, and their prediction ability will be decreased.

Recently, deep learning as a branch of machine learning has become increasingly popular. It can learn a complex data representation in the higher dimension and make a more accurate prediction than traditional machine learningbased methods. [20]. With its usefulness, deep learning has been applied to facilitate the development of computational biology. Specifically, it outperforms traditional machine learning-based algorithms in numerous drug repositioning, visual screening, and drug-target profiling [21], [22], [23], [24], [25], [26], [27], [28]. By reducing the noise, deep learning helps to extract a better drug representation and other biological data. [29], [30].

In drug response problems, deep learning is utilized to automatically learn genomic features and transcriptomic and epigenomic features of cell lines. Also, it can extract the chemical structures of drugs to predict the sensitivity of anticancer drugs. Therefore, deep learning does not need to calculate molecular features or perform feature selection, which is susceptible to errors. Various models have been proposed to address this issue. [31], [32], [33], [34]. For instance, DeepDR, tCNNS, and CDRScan can be considered as some of the original works in the field. Using a deep neural network, DeepDR predicts the half-maximal inhibitory concentrations (IC50) by building a multiple subnetwork to learn the drug representation. Besides, tCNNS uses a convolutional neural network to extract the features of drugs from SMILES string representations. However, since drug and cell line data contain redundant information, other models apply dimension reduction techniques such as autoencoder and variational autoencoder to solve this problem. Specifically, DeepDSC [33] uses deep autoencoder to extract genomics features of cell lines from gene expression data and then combine them with chemical features of compounds to predict drug responses. Other methods have been tried to solve this problem from a different perspective, such as MOLI, which is a drug-specific model, but shares the same approach of using convolution neural network as DeepDR [35]. These deep learning models generally use either strings or numerical drugs, which are not natural data presentations. Therefore, the structural information of drugs may be lost. As a result, some recent methods has represented drug structure as a graph and used graph convolutional networks to learn drug feature.

Graph convolutional networks (GCN) have been applied to learn representations of compound structures depicted as molecular graphs [36] [37]. GraphDRP [36] outperforms other string-representation-based models such as tCNNs by using GCN to represent the drugs’ graph where the edges are the bonding of atoms. There are also studies suggesting that these omics data are more informative than genomic data of cell lines [38]. Therefore, GraomicDRP [37] was recently published and shown to be the state-of-the-art method among other deep learning-based methods in drug response prediction tasks. GraomicDRP is built based on the GraphDRP model. However, instead of using only genomic data, GraomicDRP has combined different types of -omics data, namely (i.e., gene expression) and epigenomic data (i.e., methylation) to increase the performance. As a result, our work will be compared directly with GraomicDRP.

In this study, we propose GrapTransDRP (Graph Transformer for drug response prediction), a novel neural network architecture capable of extracting a better drugs representation from molecular graphs to predict drug response on cell lines. Graph Transformer was integrated with the combination of GAT-GCN to enhance the ability to predict more accurate drug responses. We also incorporated the idea of using multi-omics data from GraomicDRP. We compared our method with GraphDRP and GraomicDRP in both single-omics and multi-omics data [37] [36]. Experimental results indicate that our method performs better in root mean square error (RMSE) and Pearson correlation coefficient for all experiments. Furthermore, while both genomic and epigenomic data are presented in a binary format with 377 and 735 dimensions, respectively, transcriptomic data are continuous and normalized in the range from 0 to 1 with 17737 dimensions. Therefore, Kernel PCA was applied to reduce the dimension of into the same dimension with genomic and epigenomic data. Then our experiment indicates that the combination between Graph Transformer and GAT-GCN using all processed -omics data achieves the highest result among all possible combinations.

## 2 Graph Transformer for drug response prediction (GraTransDRP)

This study proposed the model GraTransDRP, which takes chemical information of drugs and multi-omics data of cell lines to predict response values. The proposed model is shown in Fig 1

**Fig. 1.**
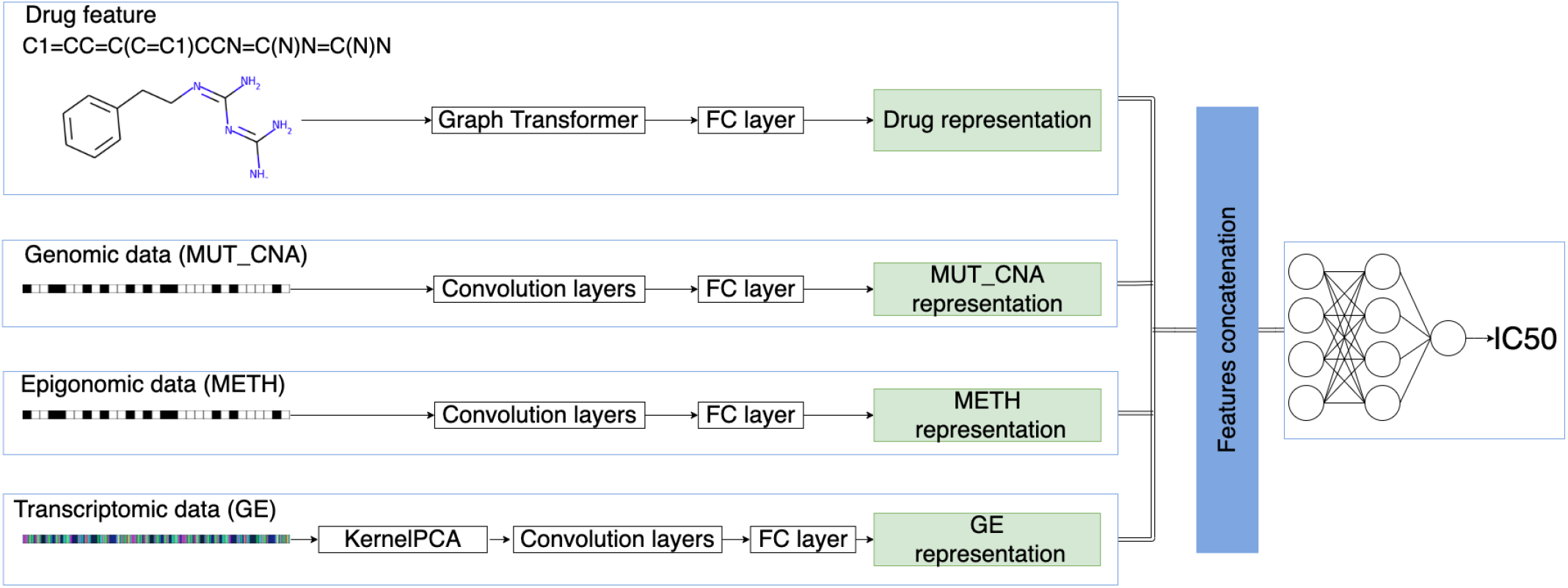
An illustration of GraTransDRP. Genomic data and epigonomic data were converted to one-hot format with a vector of 735 and 337 dimensions respectively. initially was put through KernelPCA to reduce dimension. Then 1D convolutional layers were applied three times to these features. After that, the fully connected (FC) layer was used to convert ConvNet results in 128 dimensions. Drug in SMILE string was converted to graph format. Then graph transformer were used to learn the drug’s feature. Following the graph transformer, the fully connected layer was also used to convert the result to 128 dimensions. Finally, three feature representations and drug representation were then concatenated and put through two FC layers to predict the IC50 value.

For the drug features, the drugs represented in SMILES format [39] were downloaded from PubChem [40]. Then, RDKit, an open-source chemical informatics software [41], was used to construct a molecular graph reflecting interactions between the atoms inside the drug. Atom feature design from DeepChem [42] was used to describe a node in the graph. Each node contains five atom features: atom symbol, atom degree calculated by the number of bonded neighbors and Hydrogen, the total number of Hydrogen, implicit value of the atom, and whether the atom is aromatic. These atom features constituted a multi-dimensional binary feature vector [43]. If there exists a bond among a pair of atoms, an edge is set. As a result, an indirect, binary graph with attributed nodes was built for each input SMILES string. Graph Transformer was integrated with GAT-GCN model [43] to learn the features of drugs. Following the graph neural network, a fully connected layer (FC layer) was also used to convert the result to 128 dimensions.

The genomic and epigenomic features of cell lines were represented in one-hot encoding. As for, KernalPCA was applied to reduce the dimension from 17737 to 1000. Then, 1D convolutional neural network (CNN) layers were used to learn latent features on those data. The output of each feature was put through a fully connected layer to output a 128 dimension vector of cell line representation. Finally, the 512-dimension vector, the combination of drug’s feature and cell line’s features, was put through two fully-connected layers with the number of nodes 1024 and 256, respectively, before predicting the response.

### 2.1 Graph Convolutional Networks (GCN)

Formally, a graph for a given drug *G* = (*V, E*) was stored in the form of two matrices, including feature matrix *X* and adjacency matrix *A*. *X* ∈ *R^N×F^* consists of N nodes in the graph and each node is represented by *F*-dimensional vector. *A* ∈ *R^N×N^* displays the edge connection between nodes. The original graph convolutional layer takes two matrices as input and aims to produce a node-level output with *C* features each node. The layer is defined as:

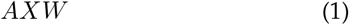

where *W* ∈ *R^F×C^* is the trainable parameter matrix. However, there are two main drawbacks. First, for every node, all feature vectors of all neighboring nodes were summed up but not the node itself. Second, matrix A was not normalized, so the multiplication with A will change the scale of the feature vector. GCN model [44] was introduced to solve these limitations by adding identity matrix to A and normalizing A. Also, it was found that symmetric normalization achieved more interesting results. The GCN layer is defined by [44] as

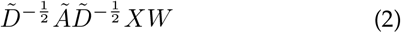

where 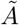 is the graph adjacency matrix with added self loop, 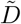 is the graph diagonal degree matrix.

### 2.2 Graph Attention Networks (GAT)

Self-attention technique has been shown to be self-sufficient for state-of-the-art-level results on machine translation [45]. Inspired by this idea, self-attention technique was used in graph convolutional network in GAT [46]. We adopted a graph attention network (GAT) in our model. The proposed GAT architecture was built by stacking a *graph attention layer*. The GAT layer took the node feature vector **x**, as input, then applied a linear transformation to every node by a weight matrix **W**. Then the *attention coefficients* is computed at every pair of nodes that the edge exists. The coefficients between node *i* and *j* were computed as

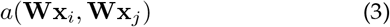

This value indicates the importance of node *j* to node *i*. These *attention coefficients* were then normalized by applying a soft-max function. Finally, the output features for each node was computed as

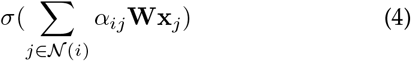

where *σ*(.) is a non-linear activation function and *α_ij_* are the normalized *attention coefficients*.

### 2.3 Combined graph neural network (GAT&GCN)

A combination of GAT [46] and GCN [44] was also proposed to learn graph features [43]. At first, the GAT layers learned to combine nodes in an attention manner, so after GAT layers, these features in each node were abstract and contained high-level information of the graph. Finally, GCN layers were used to learn convolved features and combine these abstract nodes to make final predictions.

### 2.4 Graph Transformer

Recently, Graph Attention Network (GAT) and Graph Convolution Network (GCN) have been used to learn the drug representation. These techniques are designed to learn on homogeneous graphs, while a drug’s graph can be a heterogeneous graph containing different node and link types. Also, missing connections among nodes in the graph should be taken into account when extracting the features. Graph Transformer could learn better features for a more generalized drug graph by considering these missing links. Transformer techniques have been well applied in natural language processing to solve the bottlenecks of Recurrent Neural Network (RNN). It uses multi-head attention to allow a word to attend to each other terms in a sentence. Inspired by this idea, Graph Transformer allows for the contribute of neighborhood nodes in extracting graph features. Formally, a heterogeneous graph *G* = (*V, E*) has a set of node type, 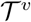, and a set of edge type 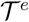. Compactly, there is an adjacency tensor *A* ∈ *R^N×N×K^*, where 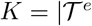, and feature matrix *X* ∈ *R^N×F^* consists of N nodes in the graph and each node is represented by *F*-dimensional vector. Also, a meta-path to predict new connections among nodes is defined as

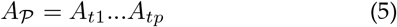

where *A_ti_* is an adjacency matrix for the i-th edge type of meta-path. For each adjacency matrix, a soft adjacency matrix Q using 1×1 convolution is chosen as

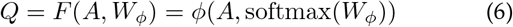

where *ϕ* is a convolution layer and *W_ϕ_* ∈ *R*^1×1×K^. Multiply these soft adjacency matrix Q together, a convex combination of new meta-paths is established. Combining with Graph Convolution network, node representations are formed as

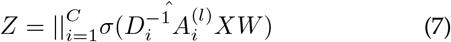

The above equation is a function of neighborhood connectivity, which is similar to the attention mechanism shown in the transformer for natural language processing. To enhance the prediction accuracy, Graph Transformer considers the positional encoding, which encodes the distance-aware information. In contrast to the natural language processing problem, the position of nodes is difficult to determine when extracting features due to the natural properties of the graph. Therefore, Graph Transformer uses Laplacian eigenvectors to address this issue. These eigenvectors are computed as

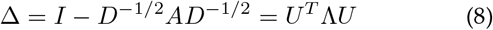

where U and Λ are eigenvectors and eigenvalues, respectively. Grah Transformer uses Laplacian eigenvectors to precompute all graphs in the dataset. In our model, we used two layers of Graph Transformer with the combination of GAT-GCN in the middle to enhance the ability to extract features and increase the prediction’s accuracy.

### 2.5 Kernel PCA

Kernel PCA is an extension of PCA [47] using kernel methods. Besides the curse, there is also a bless of dimensionality such that N points can almost always be linearly separable. While PCA is a linear method, Kernel PCA deals with a non-linear situation where it uses the kernel function to project the dataset into a higher dimensional space. Instead of working in the high dimensional space, N-by-N kernel dimensional space is created as

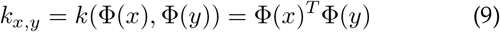

where Φ can be chosen arbitrarily, and each element in the matrix represents the similarity of one transformed data with respect to all the transformed data. Then, the same procedure as PCA on this matrix is applied. Among various kernel functions, radial basis function (RBF) is the most well-known kernel. With two samples *x* and *x′*, the kernel on two samples is calculated as

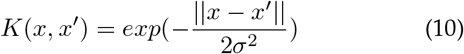

where *σ* is the width of the Gaussian distribution, the resulting feature is bounded between zero and one. It is close to one if *x* is similar to *x′* and close to zero if two data points are dissimilar. Our study chose RBF as the kernel function to reduce the dimension of gene expression data to combine with methylation and mutation data to predict drug response.

## 3 Experimental Setting

### 3.1 Datasets

CCLE [4] and GDSC [3] are extensive drug sensitivity screening projects containing not only –omics but also drug response data for anti-cancer drugs on thousands of cell lines. The –omics data includes gene expression (i.e., transcriptomic data), which indicates several RNAs transcribed from DNA and thus the amount of translated proteins in a cell. Therefore, the expression level of a gene indicates the activity level of a gene in a particular state (e.g., diseased or normal) in a cell. In addition, the –omics data also implies genomic aberrations such as mutations and copy number variations (CNVs) in genome. In this study, we also used methylation (epigenomic data), which regulates gene expression by recruiting proteins involved in gene repression or inhibiting transcription factors’ binding to DNA. Meanwhile, cell lines are cultured and treated with different doses of drugs. Therefore, we used IC50 as the drug response measure of efficiency to inhibit the vitality of cancer cells. Specifically, it indicates the amount of a particular drug needed to inhibit the biological activity by half.

GDSC is the most extensive database containing a diversity of -omics data and cell line information. In a previous study of GraomicDRP [37], there are 223 drugs tested on 948 cell lines in that database. Meanwhile, only 24 drugs were tested on 504 cell lines in CCLE. Thus, we selected GDSC for this study. Similarly, we have three types of -omics data, including Gene expression (GE), mutation and copy number aberration (MUT_CNA), and methylation (METH). In addition, a cell line was described by a binary vector, where 1 or 0 indicate whether a cell line has or has not a genomic aberration, respectively. Methylation data is also in binary form to show whether a gene is hyper-methylated or hypomethylated. Gene expression data provides a continuous expression level for genes, which is normalized between 0 and 1. The response values in terms of IC50 were also normalized in a range (0,1) as in [48]. Meanwhile, drugs were represented in canonical SMILES format [39].

### 3.2 Experimental design

In this section, the performance of our model is demonstrated through two experiments: the integration of Graph Transformer with single and multi-omics data and the effect of dimensional reduction on gene expression data. Of all cell line pairs from the GDSC database, we split it into 80% as the training set, 10% as the validation, and 10% as the test set. The validation set is used to tune the hyperparameters, and the testing set is used to evaluate the model’s generalization. Initially, we chose the values of the hyperparameters based on previous work. Then we tuned many parameters such as learning rate and batch size to achieve a better result. Detailed experiments are described below.

#### 3.2.1 The integration of Graph Transformer

This experiment aims to test the effect of the Graph Transformer model with all combinations of -omics data. Previously, GraomicDRP was tested single and multi-omics data to observe which combination achieves the highest result. In this experiment, Graph Transformer was combined with GAT-GCN and evaluated on the same procedure as before. First, the performance of the model was recorded when testing with single-omics data. Then, the model was evaluated with a combination of two and all-omics data, respectively. Finally, we compared the results to discover which variety of -omics data has the highest prediction accuracy.

#### 3.2.2 The effect of dimensional reduction on gene expression data

This experiment aims to study the effect of different dimensional reduction techniques on gene expression data to obtain the overall result. Observing the dataset, we noticed that the dimension of gene expression data is significantly larger than the dimension of mutation and methylation data. Therefore, it potentially contains redundant information and incurs noise in our prediction. We investigated this effect using dimensional reduction techniques on gene expression features, then compared the performance with original omics data and the best result in the first experiment. Specifically, PCA, KernelPCA, and Isomap were used to reduce gene expression data.

### 3.3 Performance Evaluation

Root mean square error (RMSE) and Pearson correlation coefficient (*CC_p_*) are adopted to evaluate the performance of models. RMSE measures the difference between the predicted value and the true value

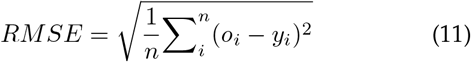

where *n* is the number of data points, *o_i_* is a ground-truth of *i^th^* sample, *y_i_* is the predicted value of *i^th^* sample.

Pearson correlation coefficient measures how strong a relationship is between two variables. *CC_p_* is defined as

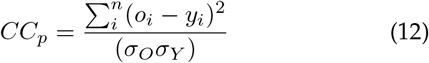

where the standard deviation of O and Y be *σ_O_*, *σ_Y_* respectively.

The lower RMSE, the better the model is. Meanwhile, the higher *CC_p_*, the better the model is.

## 4 Results and Discussion

### 4.1 The integration of Graph Transformer

Tables 2 & 3 present the prediction performance in terms of CC_p_ and RMSE for different experiments by the baseline (GraomicDRP [37]) and our proposed method (GraTrans-DRP).

**TABLE 1.**
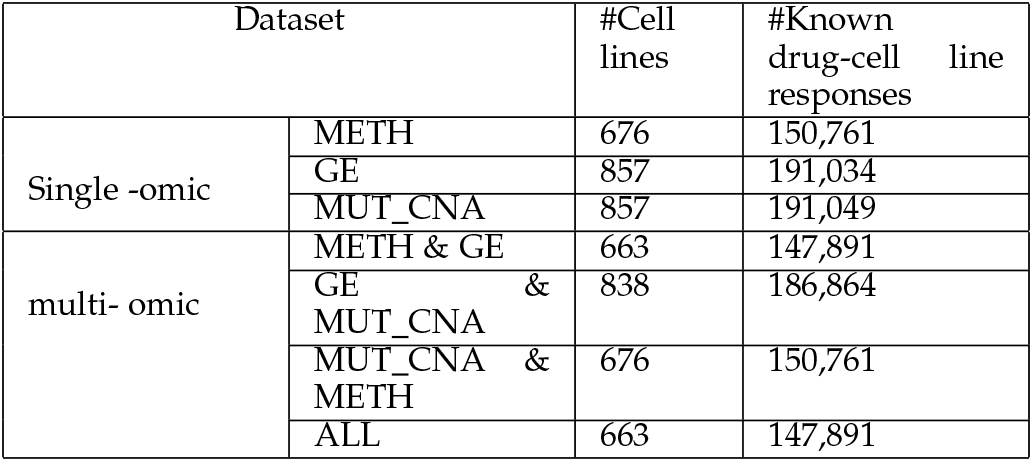
Dataset

**TABLE 2.** Performance comparison in terms of *CC_p_* and RMSE on the GDSC dataset with the integration of single-omics data. The best performance is in bold.

| Method | GraTransDRP |  | GraomicDRP |  |
| --- | --- | --- | --- | --- |
| | $CC_p$ | RMSE | $CC_p$ | RMSE |
| METH | 0.9218 | 0.0251 | 0.9104 | 0.0279 |
| GE | 0.9279 | 0.0249 | **0.9165** | **0.0259** |
| MUT_CNA | **0.9317** | **0.0238** | 0.9120 | 0.0263 |

**TABLE 3.** Performance comparison in terms of *CC_p_* and RMSE on the GDSC dataset with the integration of multi-omics data. The best performance is in bold.

| Method | GraTransDRP |  | GraomicDRP |  |
| --- | --- | --- | --- | --- |
| | $CC_p$ | RMSE | $CC_p$ | RMSE |
| METH & GE | 0.9351 | 0.0236 | **0.9310** | **0.0239** |
| GE & MUT_CNA | 0.9301 | 0.0337 | 0.9236 | 0.0246 |
| MUT_CNA & METH | **0.9353** | **0.0235** | 0.9277 | 0.0252 |
| ALL | 0.9341 | 0.0239 | 0.9295 | 0.0244 |

**TABLE 4.**
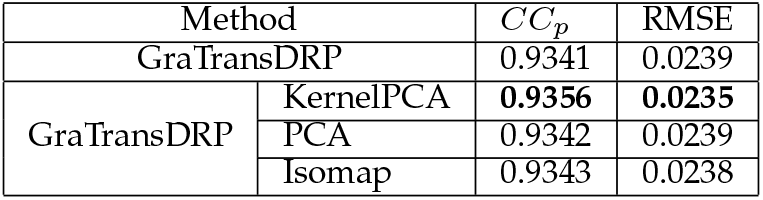
Performance comparison in terms of *CC_p_* and RMSE on the GDSC dataset with the integration of multi-omics data. The best performance is in bold.

| Method | | $CC_p$ | RMSE |
| --- | --- | --- | --- |
| GraTransDRP |  | 0.9341 | 0.0239 |
| GraTransDRP | KernelPCA | **0.9356** | **0.0235** |
|  | PCA | 0.9342 | 0.0239 |
|  | Isomap | 0.9343 | 0.0238 |

Table 2 and table 3 show the prediction performance of the models using single and multi-omics data. We observed that our proposed models, for all kinds of -omics data, achieved better RMSE and *CC_p_* than GraomicDRP. It means the Graph Transformer is more efficient than Graph Convolutional Network in extracting drugs features.

For single-omics data, the GraomicDRP’s experiment showed gene expression data has the best performance in terms of RMSE (0.0259) and *CC_p_* (0.9165). Meanwhile, our model suggested that mutation and copy number abberration data has the highest performance (0.9317 for *CC_p_* and 0.0238 for RMSE). Within each type of -omics data, we can observe that GraTransDRP outperforms GraomicDRP for *CC_p_* and RMSE. For multi-omics data, while the combination of methylation and gene expression achieves the highest performance in GraomicDRP model, GraTransDRP indicates that the combination of mutation and methylation achieves the higher result. Specifically, the *CC_p_* result of MUT_CNA & METH in GraTransDRP is 0.9353, which is higher than *CC_p_* result of METH & GE in GraomicDRP (0.9310). Similarly, we observed that the performance of GraTransDRP outperforms the performance of GraomicDRP in all combinations.

However, both models point out that combination of all three -omics data does not necessarily have the best performance. We suspected this due to redundant information introduced by new types of -omics data. Therefore, we experimented different dimensional reduction techniques on gene expression data to investigate whether the dimensionality reduction techniques improve the result.

### 4.2 The effect of dimensional reduction on gene expression data

In this experiment, we applied techniques for dimensionality reduction, namely PCA [47], Isomap [49], and KernelPCA [50]. PCA can be viewed as linear method, while KernelPCA and Isomap are better methods for non-linearity cases. We first applied these techniques on gene expression and then combined all three -omics data to use in the final model. Overall, applying dimensional reduction techniques improves the performance of the previous performance (0.9341) in terms of *CC_p_*. Among three dimensional reduction techniques, we observed that KernelPCA achieved the best performance, with 0.9356 for *CC_p_* and 0.0235 for RMSE. This experiment suggests that the gene expression data is not linearly separable in the current dimension. Instead, KernelPCA and Isomap methods project the current data into the higher dimensional space, where there exists a linearly separable hyperplane through the dataset. Since Kernel PCA has a higher experimental result, we focused on the analysis of this technique.

Graph Transformer using all -omics data with KernelPCA technique achieves the best prediction performance in terms of both RMSE and *CC_p_* among all experiments in this study. It confirms our suspicion that integrating more features is not necessary good in terms of performance as there are the redundency in omic datas. As a result, applying dimensional reduction techniques to -omics data before putting into the model improves the performance. Also, it unleashed the potential of Graph Transformer in graph representation, partly supporting the claim in [51] that Graph Transformer extracts better node features in the heterogeneous graph.

## 5 Conclusions and Discussion

In this study, we propose a novel method for drug response prediction called GraTransDRP. Our model used Graph Transformer, similar to the well-known Transformer technique in natural language processing, to extract a better drug representation. The original GAT-GCN model is combined with Graph Transformer to enhance the features of drugs and the overall drug response prediction. Instead of using only a single-omics data as in GraphDRP, we also tested our new model on multi-omics data. Using the 1D convolutional neural network, we extracted the cell line features of multi-omics data before combining them with drug representation. Also, since the dimension of is significantly larger than the other two -omics data, KernelPCA was applied to this data to reduce the noise it incorporates into the final prediction. Finally, all features were combined to predict the IC50 value. The performance of our proposed method was compared against the implementation of GraphDRP [36], and GraomicDRP [37]. The experimental results indicate GraTransDRP outperformed GraphDRP and GraomicDRP in all experiments.

In this study, we only focused on improving the data representation of the drug, which potentially increases IC50 values by using Graph Transformer. Also, in our study, we concentrated on improving drug response prediction by extracting drug features from their graph representation by GCN. Moreover, after discovering that combining three cell line elements is not necessarily equivalent to the highest performance, we apply KernelPCA to reduce the noise and obtain a better result. Finally, we used the same dataset with the GraomicDRP study to compare the relative performance, which reached the performance using single and multi-omics data. In future work, we can potentially discover other techniques to extract an even better drug representation.

---

**Tuan Nguyen** is a lecturer at National Economics University. He received his MSc degree from University of Bristol in 2019 in the area of machine learning. His current research topic includes application of deep learning in computer vision, natural language processing and bioinformatic.

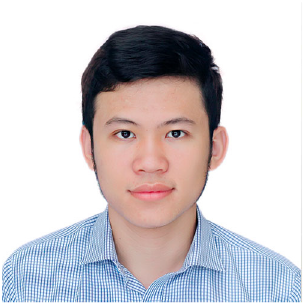

**Thang chu** is a third year student studying computer science at University of Alberta. His current research topic includes application of deep learning in biology.

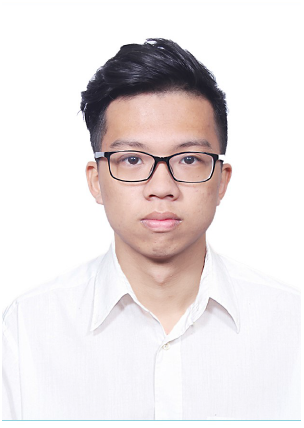

